# Environmental decomposition of cuticular hydrocarbons generates a volatile pheromone that guides insect social behavior

**DOI:** 10.1101/773937

**Authors:** Eduardo Hatano, Ayako Wada-Katsumata, Coby Schal

## Abstract

Once emitted, semiochemicals are exposed to reactive environmental factors that may alter them, thus disrupting chemical communication. Some species, however, might have adapted to detect environmentally mediated breakdown products of their natural chemicals as semiochemicals. We demonstrate that air, water vapor, and ultraviolet (UV) radiation break down unsaturated cuticular hydrocarbons (CHCs) of *Periplaneta americana* (American cockroach), resulting in the emission of volatile organic compounds (VOCs). In behavioral assays, nymphs strongly avoided aggregating in shelters exposed to the breakdown VOCs from cuticular alkenes. The three treatments (air, water vapor, UV) produced the same VOCs, but at different time-courses and ratios. Fourteen VOCs from UV-exposed CHCs elicited electrophysiological responses in nymph antennae; 10 were identified as 1-pentanol, 1-octanol, 1-nonanol, tetradecanal, acetic acid, propanoic acid, butanoic acid, pentanoic acid and hexanoic acid. When short-chain fatty acids were tested as a mix and a blend of the alcohols and aldehyde was tested as a second mix, nymphs exhibited no preference for control or treated shelters. However, nymphs avoided shelters that were exposed to VOCs from the complete 10-compound mix. Conditioned shelters (occupied by cockroaches with feces and CHCs deposited on the shelters), which are normally highly attractive to nymphs, were also avoided after UV-exposure, confirming that breakdown products from deposited metabolites, including CHCs, mediate this behavior. Our results demonstrate that common environmental and anthropogenic agents degrade CHCs into volatile semiochemicals that may serve as necromones or epideictic pheromones, mediating group formation and dissolution.

**Significance Statement:** Cuticular hydrocarbons (CHCs) cover the outer surface of insects, where they prevent water loss and serve as sex pheromones and in nest-mate recognition in social insects. Although CHCs are not volatile, they can be broken into volatile fragments by reacting with environmental agents. We demonstrate that volatile breakdown products of CHCs affect the social behavior of the American cockroach. A synthetic mix of volatiles dispersed cockroaches away from shelters, signaling an unsuitable shelter. These results highlight that some insect species have evolved communication strategies that exploit environmental and anthropogenic agents to produce bioactive compounds that mediate ecological interactions.

## Introduction

Mobile animals must assess habitat conditions to secure resources, such as shelter, food, or mates, and avoid risks, such as predators, parasites, or pathogens. Olfactory cues (semiochemicals) play a prominent role in habitat assessment because they convey information downwind before the receiver contacts the potentially hazardous source (1). For many species, aggregation pheromones – semiochemicals emanating from conspecifics – signal that a resting or feeding site may be suitable (2). On the other hand, epideictic (marking) pheromones and necromones convey risk of competition (3), hazards (4), and potentially even cannibalism (5).

Once excreted, volatile pheromones are exposed to a harsh atmosphere that may degrade the pheromone before it reaches a receiver. For instance, the honeybee Nasonov pheromone (6) is readily oxidized in air, and even plant volatile organic compounds (VOCs), like terpenes, may polymerize or oxidize (7). Thus, exposure to environmental factors that degrade semiochemicals is considered detrimental to communication.

Nonetheless, because chemical communication has evolved under the same harsh atmospheric conditions for millions of years, it is possible that insects evolved adaptive responses. Indeed, it has been shown that behaviorally inactive insect-produced chemicals are activated and mediate attraction upon exposure to environmental factors. For instance, air oxidation of the unsaturated cuticular hydrocarbons (CHC) of female yellowheaded spruce sawfly, *Pikonema alaskensis*, produced (*Z*)-10-nonadecenal which was highly attractive to males (8). Ozonolysis of CHCs of the wheat stem sawfly, *Cephus cinctus*, also produced the attractant 9-acetyloxynonanal which yielded large trap catches of males and females in the field (9). The authors speculated that attractive VOCs might be formed by natural oxidation of double-bonds of unsaturated CHCs (9). Other volatile oxidation products with similar pheromonal activity were also reported in other species (*Macrocentrus grandii* (10), *Anoplophora glabripennis* (11) and *Drosophila melanogaster* (12)). Therefore, environmentally mediated production of volatile semiochemicals from CHCs demonstrates that some species exploit seemingly harsh environmental factors in their chemical communication.

CHCs are excellent targets for their adaptive degradation by atmospheric factors into semiochemicals. CHCs are species-specific blends of aliphatic chains that cover the entire insect body surface (13, 14) and serve multiple functions, including as pheromones (15, 16). Insect CHCs are also ingested during grooming behavior, and defecated, thus accumulating on resting sites, such as shelters (17). The process of CHC degradation is similar to rancidification of lipids in food (18) and skin (19), cleaving the double-bonds in unsaturated CHCs into short-chain VOCs, such as aldehydes, ketones and fatty acids. The multiple functions of CHCs, and their loss from the cuticle through contact, grooming, and environmental degradation, may drive their relatively high turnover to maintain a fresh CHC layer and homeostasis (20).

We hypothesized that various abiotic environmental agents degrade unsaturated CHCs deposited by *Periplaneta americana* in shelters, and the resulting VOCs could then be used by conspecifics as a pheromone to assess shelter suitability for aggregating with conspecifics. *P. americana* is particularly suitable for this investigation because the major component of its CHCs is a 27-carbon diene (two double bonds) (21). We found that environmental degradation of its unsaturated CHCs generated a mix of bioactive VOCs, including fatty acids, alcohols, ketones and aldehydes, which repelled cockroaches from aggregating in shelters, but only as a complete blend. Thus, environmental conditions play a key role in chemical communication not only by disrupting chemical cues but also by activating them from behaviorally inactive organic compounds.

## Results

### I. Generation of CHC breakdown products

We first optimized the breakdown conditions of CHCs and sampling of VOCs using UV-radiation as a standard method. All methods and results (fig. S2-S4) are described in the Supporting Information. Based on these results, all subsequent experiments used borosilicate vials, positioned 15 cm from a UV-transilluminator, and exposed for up to 12 hr.

### II. Behavioral response to CHC breakdown products

We validated that the two-choice aggregation assay with *P. americana* lacked any side bias with two blank vials (fig. 1A, *p*=0.396), as well as UV-exposed vs. UV-protected blank vials (fig. 1H, *p*=0.321). Nymphs avoided the shelters associated with VOCs from CHCs exposed to air for 7 days and preferred to aggregate under blank shelters (fig. 1D, *p*<0.001) but shorter exposure to air did not generate avoidance responses (1 day exposure: fig. 1B, *p*=0.352; 3 days exposure: fig. 1C, *p*=0.597). Nymphs also avoided VOCs from CHCs exposed to water vapor for 3 days (fig. 1F, *p*<0.05) and 7 days (fig. 1G, *p*<0.001) but not for 1 day (fig. 1E, *p*=0.636).

**Figure 1.**
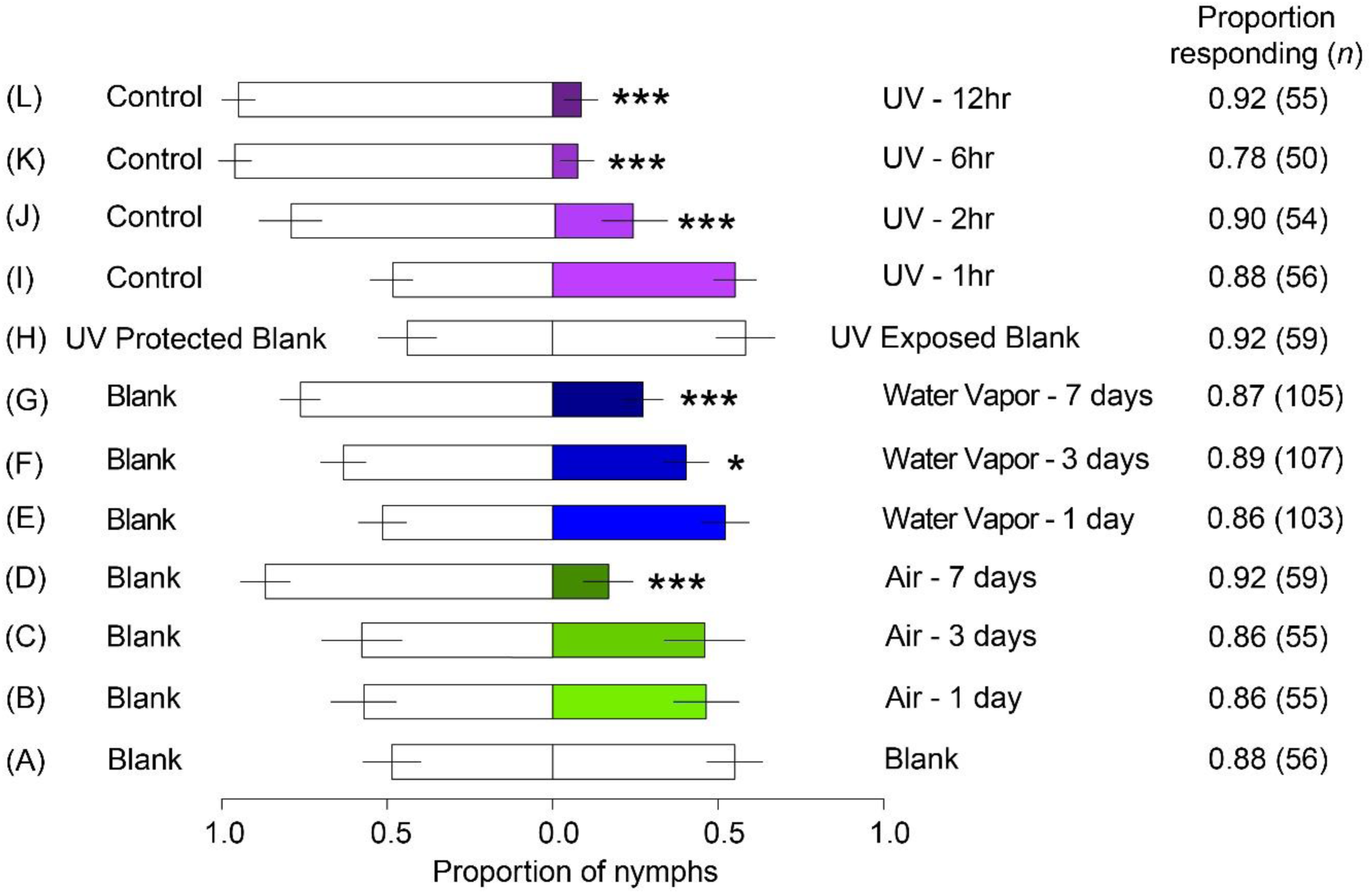
Behavioral responses of *P. americana* nymphs to breakdown VOCs of unsaturated CHCs. Groups of four second instar nymphs were tested in two-choice shelter assays in Petri dishes. The source of VOCs was a vial positioned under each shelter. Experiments were to test (A) for side bias using only vials treated with hexane (Blank); (B-D) breakdown VOCs produced when CHCs were exposed to air for 1, 3 and 7 days (Air); (E-G) breakdown VOCs produced when CHCs were exposed to water vapor for 1, 3 and 7 days (Water Vapor); (H) blank vials exposed to UV (UV-exposed blank) and protected from UV (UV-protected blank) for 12hr; and (I-L) breakdown VOCs produced when CHCs were exposed to UV-radiation for 1, 2, 6 and 12hr (UV). Bars represent mean (±SE) proportions of nymphs (lme, *p<0.05, *** p<0.001, *N* = 15-30). The proportion of tested nymphs that responded to both stimuli and respective number of nymphs (*n*) are shown next to each bar.

Exposure to UV-radiation elicited avoidance of the resulting VOCs at much shorter periods of exposure, with 1hr exposure to UV eliciting no avoidance (fig. 1I, *p*=0.476), but 2, 6 and 12hr of exposure resulting in strong avoidance (fig. 1J-L; *p*<0.001).

### III. Analysis of VOCs

When unsaturated CHCs were exposed to air, no VOCs were detected on day 1 compared to the respective control vial (fig. 2A-I). Small amounts of VOCs appeared after 3 days (fig. 2A-II) and more compounds at higher amounts appeared after 7 days (fig. 2A-III). Unsaturated CHCs exposed to water vapor followed a similar pattern, but the ratios of compounds were different between the two treatments (fig. 2A-IV, V, VI). In contrast, exposure of alkenes to UV-radiation produced VOCs within 1-6hr and large amounts of VOCs were generated after 12hr of UV-exposure (fig. 2VII-IX); notably, unsaturated CHCs that were protected from UV did not produce VOCs (fig. 2VII-IX) nor did clean borosilicate vials exposed to UV-radiation (fig. S5). We calculated correlation coefficients for each treatment pairing based on the relative abundance of GC peaks, and significant correlations are displayed in a correlation matrix (fig. S6). The VOCs from exposure of unsaturated CHCs to air or water vapor for 7 days, or to UV for 6 or 12hr were significantly positively correlated (fig. S6). At these time points, the VOC compositions were similar, but their concentrations and ratios differed (fig. 2A-III, VI, VIII and IX); yet, all three methods elicited the same behavioral responses in *P. americana* nymphs. Of the three methods, however, UV produced the highest amounts of VOCs in the shortest time, including all the VOCs produced under the effects of water vapor and air. Therefore, UV-exposure was used for all subsequent analyses, including GC-EAD and GC-MS.

**Figure 2.**
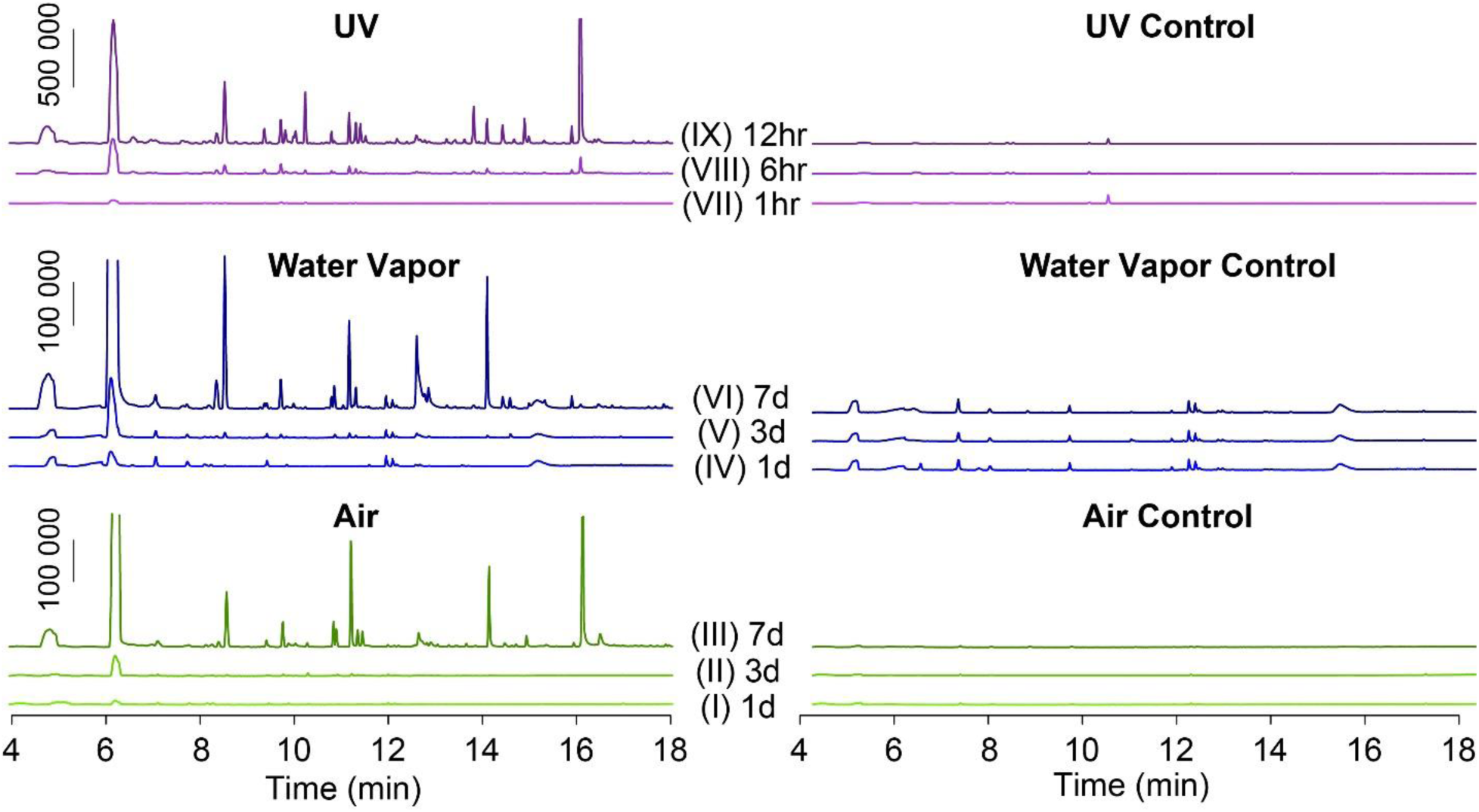
Time-courses of production of VOCs from degradation of *P. americana* unsaturated CHCs. CHCs were exposed to air, water vapor or UV-radiation. Total ion chromatograms of breakdown VOCs of unsaturated CHCs. Left: VOCs emitted from CHCs exposed to air (I-III) or water vapor (humidified air) (IV-VI) for 1, 3 and 7 days, or UV-radiation (VII-IX) for 1, 6 and 12hr. Right: chromatograms of the respective controls for air (blank vials, I-III), water vapor (blank vials, IV-VI) and UV-radiation (CHCs protected from UV-radiation: VII-IX). VOCs were sampled by SPME.

### IV. Identification of bioactive VOCs

Active VOCs from UV-exposed unsaturated CHCs were recognized using gas chromatography coupled to electroantennographic detection (GC-EAD) with nymph antennae. Fourteen compounds elicited antennal responses and 10 of these were identified by mass spectrometry (fig. 3). Consistent with the behavioral results, bioactive compounds gradually accumulated over time of UV-exposure of unsaturated CHCs. Among these, five compounds were short-chain fatty acids (acetic, propanoic, butanoic, pentanoic and hexanoic acids) which comprised 68% of the total mass of volatiles. The other compounds (26%) included tetradecanal, 1-pentanol, 1-octanol, 1-nonanol and 2-nonanone. Unsaturated CHCs that were not exposed to UV did not produce any EAD-active compounds (fig. S7).

**Figure 3.**
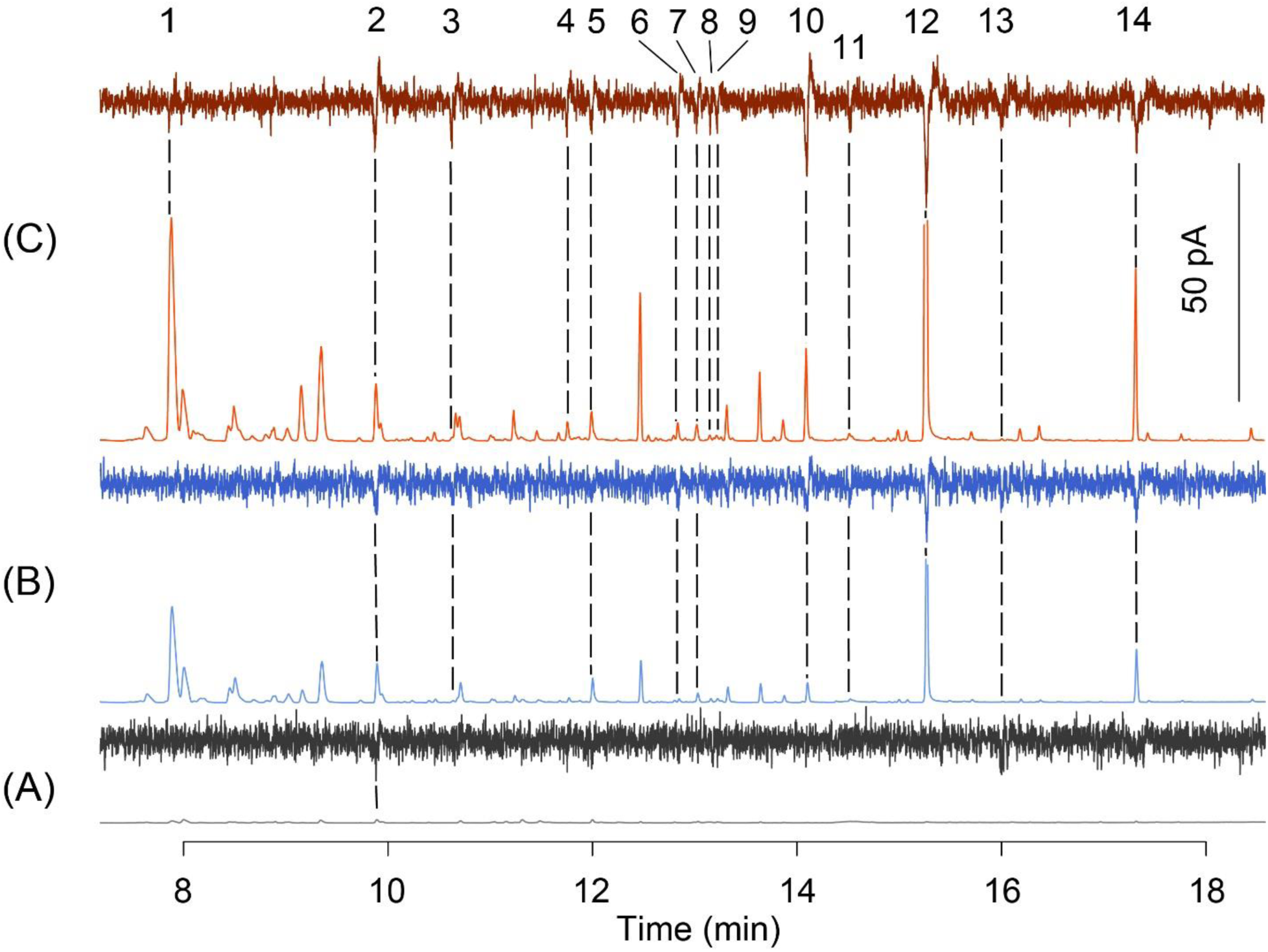
Antennal electrophysiological responses of second instar *P. americana* nymphs to volatile compounds from the UV-mediated decomposition of unsaturated CHCs for (A) 1hr, (B) 6hr and (C) 12hr. In each panel, the bottom (upwards going peaks) shows the FID output chromatogram, and the top (downward going EAD responses) shows the antennal responses; larger downward going responses indicate greater response to the corresponding FID peak (dashed lines connect FID peaks and EAD responses). Antennal response is represented by the median of EAD recordings (*N*=6-9). Compounds: 1-pentanol (1), 2-nonanone (2), acetic acid (3), propanoic acid (4), 1-octanol (5), butanoic acid (6), unknown (7), unknown (8), 1-nonanol (9), pentanoic acid (10), unknown (11), hexanoic acid (12), tetradecanal (13), unknown (14). Scale bar for FID chromatograms: 50 pA.

### V. Behavioral response to identified compounds

We prepared two synthetic mixes from the identified EAD-active compounds: one mix contained the five fatty acids, and the other mix contained the remaining five compounds (alcohols, ketone and aldehyde). These compounds were carefully mixed to reproduce the concentrations observed in the VOCs from CHC degradation. Bioassays with hexane demonstrated that cockroaches did not prefer either shelter position in the two-choice arena, that is, there was no side bias (fig. 4A, *p*=0.403). Neither mix alone affected the choice of nymphs for shelters (fig. 4B and C; Fatty Acids Mix: *p*=0.138; Other VOCs Mix: *p*=0.359). However, when both mixes were combined, cockroaches avoided the shelters positioned over the combined mixes and aggregated under the control shelters (fig. 4C, *p*<0.001), as they did in response to the natural VOCs of CHC degradation. In order to assess the effect of quantitative differences between the two blends, the concentration of the Fatty Acids mix and the Other VOCs Mix was increased 10% and 90%, respectively, to match the total mass of all the identified bioactive volatile products. We found no difference in the aggregation of cockroaches under shelters in response to the high concentration of the Fatty Acids Mix (fig. 4D; *p*=0.076). In contrast, the high concentration of the Other VOCs Mix (alcohols, ketone, aldehyde) elicited significant aggregation in the shelter above this mix (fig. 4E, *p*<0.001).

**Figure 4.**
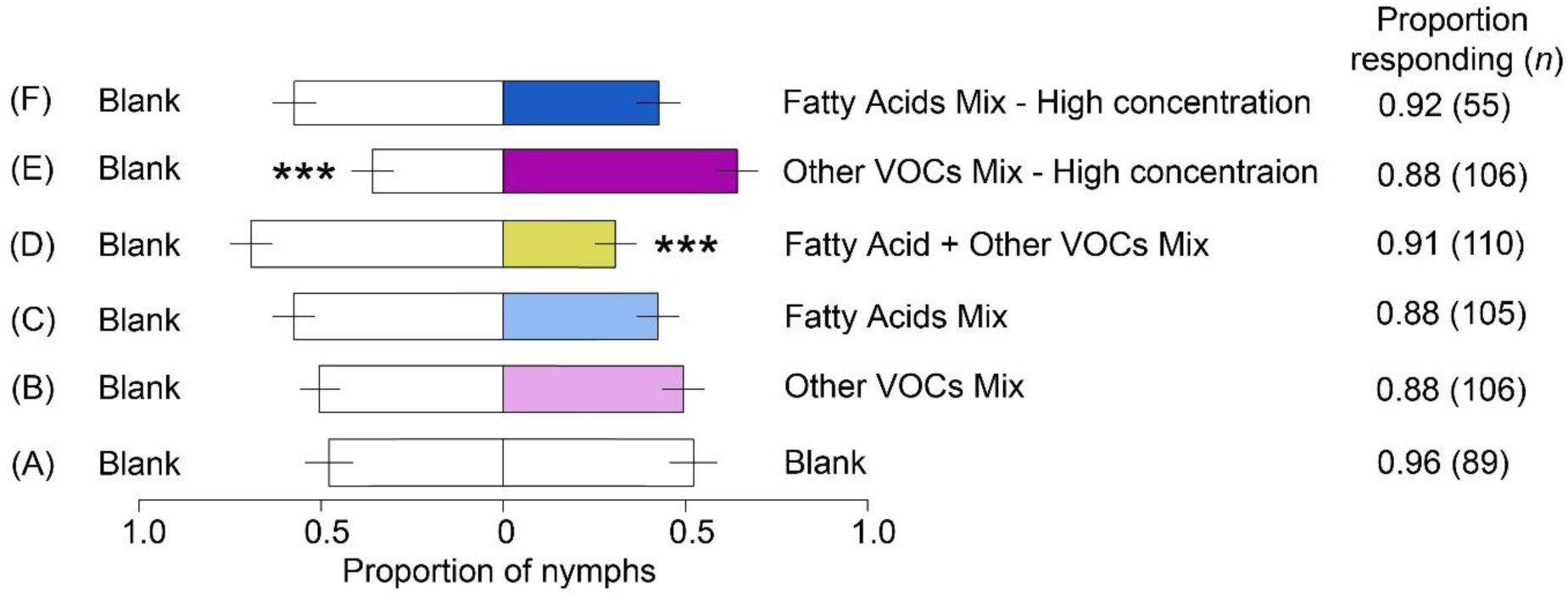
Behavioral responses of *P. americana* nymphs to synthetic mixes of EAD-active CHC breakdown VOCs. Groups of four second instar nymphs were tested in two-choice shelter assays and nymphs on each shelter were counted after 20 min. Experiments were to test (A) for side bias using vials treated with hexane control (Blank); (B) the effect of a mix of 1-pentanol, 2-nonanone, 1-octanol, 1-nonanol and tetradecanal (Other VOCs); (C) a mix of acetic, propanoic, butanoic, pentanoic, and hexanoic acids (Fatty Acids); and (D) a combination of both mixes (Acids+Other VOCs). In order to compensate for the higher concentration of fatty acids in the total VOCs, the (E) Other VOCs Mix and (F) Fatty Acids Mix were tested at concentrations matching those of the total VOCs. Bars represent mean (±SE) proportions of nymphs (lme, *** p<0.001, *N* = 15-30).

### VI. Behavioral responses to cockroach-conditioned shelters

Filter papers were conditioned by cockroaches, a procedure that results in the deposition of fecal and oral secretions and cuticular lipids on the papers. Exposure of clean shelters to UV did not affect the aggregation responses, as evidenced by lack of preference for UV-exposed vs. UV-protected clean filter papers (fig. 5A-I, *p*=0.78). As expected, the odor from conditioned shelters that were protected from UV were significantly more attractive to cockroaches than clean shelters treated in the same manner (fig. 5A-II, *p*<0.001). However, this preference was reversed when both types of shelters were exposed to UV (fig. 5A-III, *p*<0.001).

**Figure 5.**
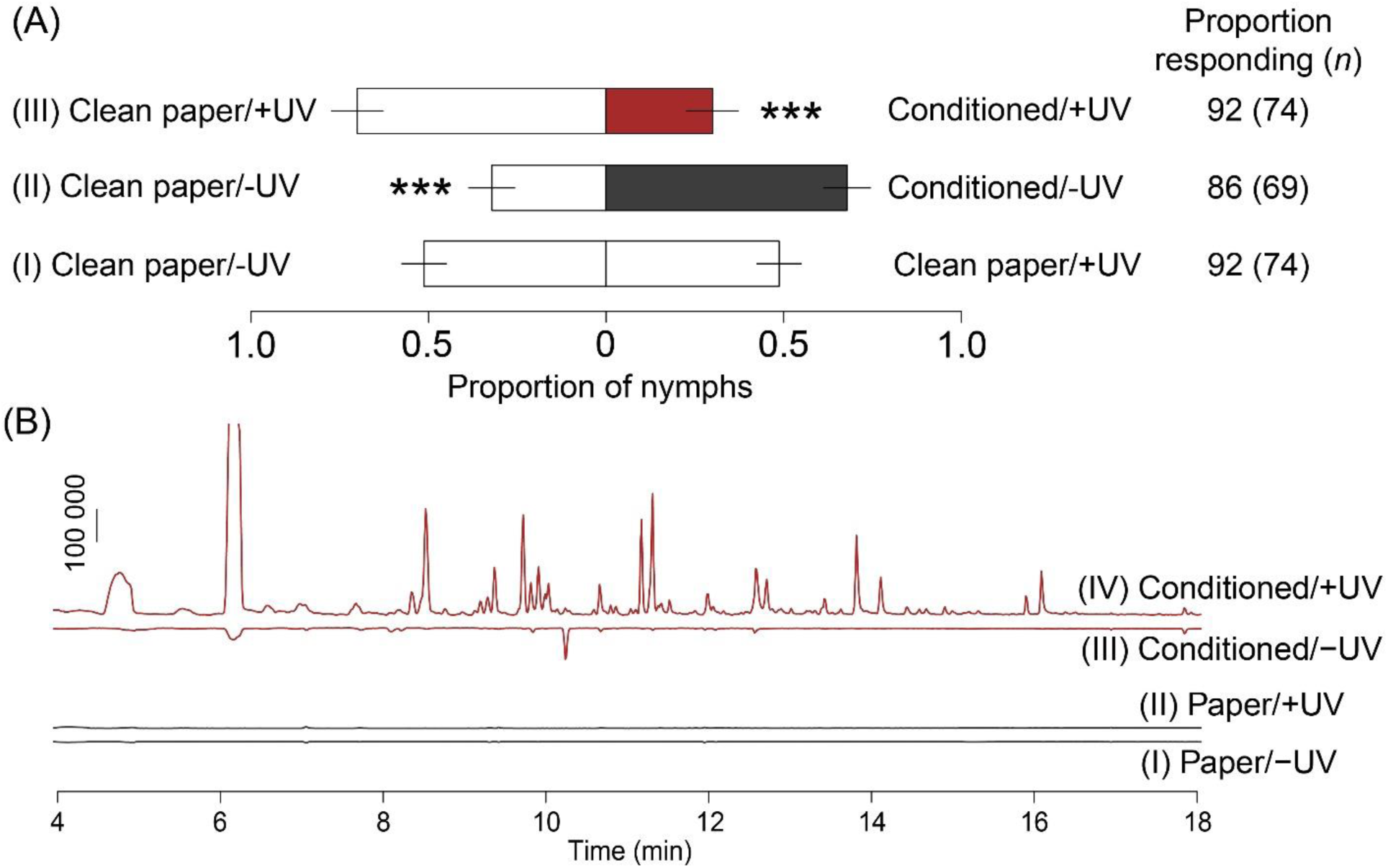
Effect of UV-radiation on CHCs deposited on conditioned filter papers. Pieces of conditioned or clean filter papers were placed inside vials and exposed to UV-radiation for 12hr. (A) Aggregation of *P. americana* nymphs on cockroach-conditioned shelters. Groups of four second instar nymphs were tested in two-choice shelter assays and nymphs on each shelter were counted after 20 min. The behavioral assays tested the effect of (I) clean containers using clean filter paper protected from UV (Clean paper / −UV) and exposed to UV (Clean paper / +UV); (II) filter papers conditioned by adults and protected from UV (Conditioned paper / −UV); and (III) filter papers conditioned by adults and exposed to UV (Conditioned paper / +UV). Bars represent mean (±SE) proportion of nymphs (lme, **p<0.01, *N* = 15-30). (B) Total ion chromatograms of odorants collected from (I) clean filter paper protected from UV and (II) exposed to UV-radiation, (III) shelters conditioned with cockroaches protected from UV and (IV) exposed to UV-radiation. Chromatograms of treatments protected from UV-radiation are facing downward, while chromatograms exposed to UV-radiation are facing upward. Scale bar: 100 000 abundance.

### VII. Accumulation and degradation of CHCs on shelters and formation of volatile products

The chemical profiles from UV-exposed and UV-protected clean shelters were similar, with little VOCs detected in either treatment (fig. 5B-I and II). Headspace analysis with solid phase microextraction (SPME) confirmed that conditioned shelters exposed to UV produced more and different VOCs than UV-protected shelters (fig. 5B-III and IV). Furthermore, the odor profile from UV-exposed conditioned papers was strikingly similar to the profile of UV-exposed unsaturated CHCs, which, as shown before, also differed from the VOCs emanating from UV-protected CHCs (fig. S8A).

GC analysis of shelter extracts showed that UV-exposed shelters contained less of the main CHC, 6,9-heptacosadiene, than UV-protected shelters (fig. S8B, *p*<0.05), while amounts of the main saturated CHCs, *n*-pentacosane and 3-methylpentacosane, did not differ (fig. S8B, *p*=0.761 and *p*=0.937, respectively).

## Discussion

The majority of insect semiochemicals have been collected and tested under controlled environmental conditions to avoid contamination and decomposition, which may result in loss of bioactivity. Natural field conditions abound with oxidizing agents (e.g., oxygen, ozone, water vapor, light, and UV-radiation), which are often unavoidable, and can modify the emitted semiochemicals. These agents may significantly reduce the life-span of semiochemicals, disrupting chemical communication between organisms (6). However, insects have evolved and perfected chemical communication mechanisms for millions of years under these harsh environmental conditions, and it is reasonable to propose that some insects, particularly those adapted to anthropogenic environments, exploit these environmental agents to perform the last step of semiochemical production. Under this hypothesis, insects produce pro-pheromones (or more generally, pro-semiochemicals), chemicals whose decomposition products under the influence of environmental agents serve as pheromones (or more broadly, semiochemicals).

The most prominent potential pro-pheromones that are readily available for interaction with atmospheric agents are CHCs. These long-chain compounds are ubiquitous in insects, occur in relatively large amounts and in species-specific blends, serve multiple unrelated physiological and behavioral functions, and are maintained at homeostatic levels through regulated biosynthesis, deposition on the cuticle, and grooming (22, 23). Some insect species contain a mix of saturated, unsaturated and methyl-branched CHCs, while others display only saturated CHCs (13). Notwithstanding, the CHC profile of *P. americana* is quite unique among insects in general (cf *Drosophila melanogaster*) and even closely related *Periplaneta* species, which contain either no alkenes or a single unsaturated CHC (24). *Periplaneta americana* CHCs include several alkenes that constitute >70% of total CHCs, and >68% is represented by a single alkadiene, (*Z,Z*)-6,9-heptacosadiene (21, 24). A review by Lockey (13) concluded that <50% of 119 insect species contained unsaturated CHCs, and 36 species had alkadienes and/or alkatrienes, but most in small amounts. It is important to note, however, that alkenes are often obscured by alkanes in GC analysis, so the representation of alkenes in insects may be underestimated.

The CHCs of *P. americana* are thought to mediate species-specific aggregation (24), but it is unknown why *P. americana* evolved such a high proportion of a single alkadiene. We suggest that the prominence of 6,9-heptacosadiene in all stages of *P. americana* make it a likely pro-pheromone for producing volatiles via environmentally mediated decomposition. Its presence in nymphs, males and females (24) would indicate that volatile reaction products might serve as pheromones perceived by all developmental stages. We focused on early instar nymphs because they exhibit strong gregarious behavior and response to semiochemicals (25), as in other cockroaches (26). In this study, we demonstrated that atmospheric agents degrade *P. americana* CHCs, releasing volatile semiochemicals that repel nymphs from settling in a shelter exposed to these VOCs. These pheromones might serve to identify old-colonized shelters (epideictic) or as necromones.

Several metabolites from *P. americana* that were thought to be intrinsically biosynthesized have been identified as aggregation pheromones (27), sex pheromone (28), and necromones (29). We found that VOCs for this species were produced from the interaction of CHCs with air, water and UV-radiation, and all elicited behavioral responses. It is unlikely that *P. americana* would be exposed to UV, because it is nocturnal and it inhabits UV-protected but chemically complex environments such as sewers, bat guano in caves, and dump sites (30). These habitats contain high concentrations of numerous chemical agents (e.g., ammonia, hydrogen sulfide, acids) (31) that could interact with CHCs to form various VOCs, including the ones found in our study. Therefore, UV-radiation was used here as a model environmental agent to effect the timely decomposition of CHCs and facilitate exploration of the function of the reaction products. Because the reaction of CHCs with air and water vapor produced the same behavioral responses as to UV-produced volatiles, the results of this proof-of-concept study should compel future studies of the effects of various ecologically relevant environmental factors, alone and in combinations, on semiochemical production.

The production of insect attractants via oxidation of long-chain CHCs was demonstrated for *P. alaskensis* (8), *M. grandii* (10), *A. glabripennis* (11), *C. cinctus* (9) and *D. melanogaster* (12). Other unsaturated metabolites may also serve as precursors to volatile cues, such as tetracosyl acetate from *Diaphorina citri* (32). Most of these studies focused on the effect of a single component within the complex VOCs (but see Wickman et al (11)) and on the effect of oxidation in air or with synthetic reagents. However, a wide range of VOCs can be formed from lipid degradation simultaneously, as multiple factors affect the process (33), such as the type of precursors (e.g., chain length, number and position of double and triple bonds), environmental factors (e.g., oxidant, temperature, radiation, metal ions) and period of exposure. While the formation of some oxidation products from unsaturated hydrocarbons can be relatively straightforward (33, 34), it may still yield multiple unpredicted components. For instance, ozonolysis of the hydrocarbon (*R*)-(+)-limonene produces approximately 1200 different compounds, among which at least 75 were of low molecular weight (35). Whereas single compounds within a VOC blend might be potential pheromones, it is essential to test the complete blend of VOCs to identify their ecological function, because olfactory coding and behavioral responses to odor blends are not the same as to individual components of the blend (36). In our studies, the composition of *P. americana* CHC decomposition products varied with degradation agent and exposure period, suggesting that the quality and quantity of environmentally generated semiochemical blends are complex and shaped by abiotic habitat conditions.

Fatty acids trigger different behavioral responses in different species. Some medium-chain fatty acids, like linoleic and oleic acids, act as necromones for *Apis mellifera* (37), *P. americana* (29), termites (38) and isopods (4), repelling them from sites with dead conspecifics to avoid potential risk of predation or pathogenesis. However, short- and long-chain fatty acids from feces of *Blattella germanica*, some of which were microbial mediated (39), elicited aggregation response (40). Here, *P. americana* CHC reaction products included compounds of different chemical classes, but short-chain fatty acids were highly predominant in quantity. Given their prominence in the emitted VOCs, and the importance of fatty acids in cockroach behavior, we examined the role of these fatty acids and the remaining VOCs separately. Our results suggest that the fatty acids alone are not responsible for repellence, and either a subset or all components in both mixes are required to elicit repellence. This observation is consistent with studies showing that combinations of components may elicit stronger behavioral responses than individual components from insects, including cockroaches (40). Furthermore, we were unable to elicit the shelter avoidance response by increasing the concentration of fatty acids, confirming that the pheromone is defined primarily by the odorant composition of the mix and secondarily by its concentration. Conversely, increasing the concentration of the Other VOCs mix (i.e., alcohols, ketone and aldehyde) elicited attraction of nymphs to the shelter that received this treatment. This unexpected result suggests that if environmental conditions favored the formation of these non-fatty acid compounds from CHCs, attraction and aggregation might be mediated in *P. americana*, rather than repellence. Hence, the formation of VOCs – and potentially semiochemicals – might be plastic, not only depending on the type of CHCs as substrates, but also varying with the prevalent environmental conditions.

The majority of the investigated cockroach species are gregarious, living in groups for long periods with few individuals dispersing to distant areas (41, 42). Recognition of their own group is mediated through chemical cues emitted directly from the cockroaches and from organic materials deposited on or near the refugium, like volatiles from feces and associated microbes (39), and contact metabolites, like CHCs (but see Hamilton et al. (17)). We hypothesize that as feces and CHCs accumulate on shelters and react with environmental factors, VOCs signal colony conditions, such as size, demographic composition and health. We observed that cockroach-conditioned shelters that were initially attractive to nymphs were later avoided after exposure to UV-radiation, which triggered changes in the VOCs, largely through degradation of cuticular alkenes. We attribute the change in VOCs to an “ageing” effect on the accumulated CHCs imposed by environmental agents. Although early developmental stages of *P. americana* tend to readily form aggregations compared to later nymphal stages and adults, they are also more susceptible to injury and cannibalism in dense aggregations (25). Thus, it is plausible that early stage nymphs benefit most from assessing the chemical cues that emanate from refugia and increase their survival rate by avoiding or dispersing from aggregations that impose high risk of mortality.

In this report we used *P. americana* and UV-radiation as a proof-of-concept model for UV-mediated production of semiochemicals from unsaturated CHCs. We recognize that *P. americana* would rarely be exposed to UV, and instead other environmental factors would cause the CHC decomposition. Other insect species, however, might display behavioral adaptations to prevent or facilitate UV-exposure and VOC production. Basking in many diurnal insects, for example, might serve not only the recognized functions of thermoregulation and intraspecific display, but it might facilitate the generation of semiochemicals from CHCs. Thus, laboratory-based chemistry studies and bioassays should be integrated with behavioral observations of insects under field conditions.

## Conclusion

Insects integrate and coordinate morphological, physiological and behavioral adaptations to optimize chemical communication, and environmental decomposition of CHCs to produce semiochemicals is emerging as an important strategy. Insects are constantly exposed to abiotic environmental factors which they might exploit by selecting habitats or periods of activity that maximize or minimize interactions with these factors. Here, we demonstrated that abiotic environmental factors – air, water vapor and UV-radiation – serve as activating agents of semiochemicals for *P. americana* through reactions with CHCs. These CHC decomposition products repelled cockroaches from refugia and disrupted their aggregation. Previous studies with other species demonstrated that volatile products attract conspecifics indicating that environmental decomposition of CHCs can produce semiochemicals that are involved in different ecological interactions. Although few species have been investigated to date, they represent diverse taxonomic groups (i.e., Lepidoptera, Coleoptera, Hymenoptera, Diptera, Hemiptera and Blattodea), leading us to speculate that this phenomenon might be a widespread adaptation among insects. Furthermore, the same strategy may be employed by other organisms, as suggested for plants (43), but yet to be tested. A range of behaviors may also contribute to the environmentally induced production of volatile semiochemicals from CHCs. Grooming behavior, for example, may remove damaged “old” CHCs and distribute fresh CHCs over the insect surface (34). Together with continuous production and deposition of CHCs on the epicuticle, grooming ensures a fresh and consistent layer of CHCs on the epicuticle. For diurnal insects, basking may enhance semiochemical production by exposing CHCs to solar radiation or heat. In addition, the epicuticle of most insects contains other lipids (e.g., triglycerides and fatty acids), albeit usually in small amounts, that can become volatile cues. Therefore, studies of the adaptive role of environmental factors in producing volatile semiochemicals should have multiple dimensions, including assessing the chemical abiotic complexity of habitats, especially at times when insects are active; elucidating the matrix of metabolites deposited on the cuticular surface and in refugia because these metabolites may serve as pro-semiochemicals; observing behaviors that may promote CHC decomposition under controlled and natural conditions; and applying multiple bioassay paradigms to elucidate the functions of VOCs produced from CHCs decomposition. Finally, future studies should also focus on comparing closely related and evolutionarily divergent species that occupy habitats with different environmental agents.

## Materials and methods

### *P. americana* rearing

*P. americana* colonies were maintained at 27 °C under a 12:12 light–dark photoperiod in plastic containers with access to water, dry LabDiet rodent chow (Purina No. 5001; PMI Nutrition International, St. Louis, MO), and molded fiber egg cartons as shelters.

### CHC extract

Fifth instar *P. americana* nymphs were placed in plastic boxes (25 × 8 × 8cm). Within 24hr after molting to adults, 10 females were reared in another box for 7 days and frozen. Females were immersed in 200ml hexanes (Fisher Scientific, Fair Lawn, NJ, USA) in a clean beaker (500ml), gently swirled for 2min and the extract was transferred to another beaker. This procedure was repeated twice more, and extracts were concentrated with a rotary evaporator (Büchi, Flawil, Switzerland).

CHCs were purified using silica column chromatography (ø=2cm; silica gel: 7 g, 70-230 mesh, EM Science, Gibbstown, NJ). CHCs were eluted with 31.5 ml hexane, and concentrated with a rotary evaporator. Silver nitrate (10%) in silica gel (230 mesh, Sigma-Aldrich, St. Louis, MO) was used to separate saturated from unsaturated CHCs by eluting the column with 31 ml hexane followed by 62 ml hexane/acetone (4:1). Fractions were confirmed using gas chromatography (see Supporting Information), and fractions containing the unsaturated CHCs were concentrated under a N2 stream and suspended in hexane to yield a concentration of 1 cockroach equivalent per ml. Solutions were stored in a −30 °C freezer.

### CHC degradation and sampling of VOCs

A 50 μl aliquot from the unsaturated CHCs solution was transferred to a clear borosilicate vial (12 × 32 mm, National, Rockwood, TN, USA) that was placed in a horizontal position and hexane was allowed to evaporate for 20 min at room temperature. Vials were flushed with a stream of N_2_ for 5 min followed by medical grade air (Airgas National Welders, Radnor, PA) for 10 seconds.

CHCs were exposed to different agents to assess the production of odorants. For air treatment, the vials were immediately capped (polypropylene cap, PTFE/silicone septa, Thermo Scientific, Germany) after flushing with medical grade air. For water vapor treatment, 4 μl of HPLC grade water (Fisher Scientific, Fair Lawn, NJ, USA) was added to each vial and the vials were capped. This volume was sufficient to form condensation in the vial within 24hr. Controls were empty clean vials that were treated as the treatment vials. Vials were kept inside a fiberboard box at room temperature for 1, 3 or 7 days.

Vials for UV-exposure were capped and placed 15 cm under a transilluminator for 1, 6 or 12hr. Caps were covered with aluminum foil to protect them from degradation under UV. Control vials also contained CHCs and were placed under the transilluminator, but they were fully covered with aluminum foil. A box fan was placed in front of the transilluminator to prevent excessive heating of the vials.

After exposing CHCs to the different agents, odorants were collected with SPME (50/30 μm, DVB/Carboxen/PDMS, 2 cm fiber length, Supelco, Bellefonte, PA) by exposing the fiber for 30 min to the interior of capped vials. Odorants were then analyzed by either GC–EAD or GC-mass spectrometry (GC-MS).

### Electrophysiology

Antennal responses of 2^nd^ instar *P. americana* nymphs to volatile collections from CHC breakdown products were studied by GC–EAD. An antenna was ablated at the base and inserted into a reference glass electrode filled with Ringer, while the recording electrode was connected to the cut tip of the antenna and connected to a custom made amplifier (44). The amplifier was connected to a G3456-60010 AIB board in a 7890 GC (Agilent Technologies, Palo Alto, CA, USA) which synchronized the outputs of the FID and EAD.

The GC was equipped and operated according to details in Supporting Information. The effluent capillary for the EAD passed through a water-cooled odor delivery tube (30 cm × 8 mm) set at 19 °C, where it was mixed with humidified medical grade air (80 ml.min^-1^). The insect antenna was positioned 0.5 cm from the outlet of the odor delivery tube.

### Identification of bioactive compounds

Volatile collections were analyzed on a GC-MS (6890 GC and 5975 MS, Agilent Technologies, Palo Alto, CA, USA) and operated according to details in Supporting Information. Compounds were identified based on Kovats indices, electron ionization mass spectra and comparison with authentic synthetic standards.

### Preparation of synthetic mixtures

Synthetic compounds were systematically mixed to reproduce the amount of odorants emitted from cuticular alkenes when exposed to UV light for 12hr. Acetic, propanoic, butanoic, pentanoic and hexanoic acids (Supelco, Bellefonte, PA) were mixed in hexane at concentrations of 760, 38, 38, 38, 760 ng.μl^-1^, respectively, and henceforth called Fatty Acids Mix. The remaining compounds, 1-pentanol, 1-octanol, 1-nonanol (Supelco, Bellefonte, PA) and 2-nonanone (Sigma-Aldrich, St. Louis, MO) were mixed as 1225, 5.5 and 7.5 and 22.5 ng.μl^-1^, respectively, and tetradecanal (Supelco, Bellefonte, PA) was prepared as a separate solution (100 ng.μl^-1^), and henceforth called Other VOCs Mix.

For behavioral assays, 4 μl of each mix was applied to a piece of filter paper (2 × 2 cm). After 10 min drying at room temperature, each filter paper was transferred to a 2 ml borosilicate vial and tested. Emission rate of VOCs from filter papers was assessed by sampling the headspace of capped vials with SPME and analyzed by GC using the same procedure described for analyzing VOCs from CHC degradation.

### Behavioral assays

All assays were conducted in a walk-in chamber (40% RH, 27 ^o^C), 1hr after the beginning of the scotophase under red light. A two-choice arena was constructed of a polystyrene petri dish (ø=8.5; fig. S1, details in Supporting Information) to which two equidistant borosilicate vials containing test materials were inserted to the base through openings covered with a metal screen. Aluminum foil that wrapped control vials was removed before it was inserted into the arena. A shelter made of a “M” shaped piece of filter paper (2.5 × 2.5 cm) was placed on top of each opening. Four 2^nd^ instar nymphs were released and the dish was covered. After 20 min, the numbers of nymphs resting on each shelter were counted. The positions of test materials were randomized to avoid side bias.

### Conditioning of shelters

Filter papers (15 × 22 cm) were cleaned with acetone (Sigma-Aldrich), and conditioned by 15 female *P. americana* as described in the Supporting Information.

### Statistical analysis

All statistical analyses were performed in R (version 3.5.1) (45). Behavioral responses of nymphs under each shelter were analyzed as proportions by linear mixed effects models (lme) using the package “lme4” (46) with positions of arenas and experiment blocks (experimental days) as random effects. All test statistics are presented in Table S1.

## Acknowledgements

We thank Rick Santangelo and Brandy Simmons for maintaining the insect colonies, and Judy Elson for tests in the Xenon-weather-ometer. This work was supported in part by the United States National Science Foundation (IOS-1557864) and the Blanton J. Whitmire endowment at North Carolina State University.

